# Enhanced immunogenicity of a *Clostridioides difficile* TcdB vaccine adjuvanted with a synthetic dual-TLR ligand adjuvant

**DOI:** 10.1101/2024.11.25.625229

**Authors:** Farha Naz, Nicholas Hagspiel, Feifan Xu, Brandon Thompson, G. Brett Moreau, Mary Young, Joel Herbein, Christopher B Fox, William A Petri, Mayuresh M Abhyankar

## Abstract

We report a comprehensive evaluation of the toxin B (TcdB) vaccine adjuvanted with a dual Toll-like receptor ligand liposome adjuvant for *Clostridioides difficile* infection (CDI). The vaccine completely protected mice from a lethal infection. Compared to alum adjuvanted TcdB, it generated functionally superior systemic antibodies and supported strong memory B cell and gut IgA responses. This pharmaceutically acceptable adjuvant platform holds promise for developing a next-generation CDI vaccine.

---

*Clostridioides difficile* is a Gram-positive, spore-forming, anaerobic bacterium responsible for antibiotic-associated severe diarrhea, especially in healthcare settings^1,2^. Most of the virulent strains produce two major exotoxins, toxin A (TcdA) and toxin B (TcdB), which cause CDI by disrupting the actin cytoskeleton of host cells^3^. At least 8 TcdB subtypes have been identified and virulent TcdA^−^TcdB^+^ strains have been clinically isolated^4^. TcdB alone is able to induce a full spectrum of disease and appears to be the most dominant virulence factor^4,5^.

There is no vaccine against CDI. Recent “Cdiffense” (Sanofi-Pasteur) and “CLOVER” (Pfizer) Phase-3 trials using alum adjuvanted vaccines failed to offer any significant protection in the vaccinated group over the control group^6,7^. The reasons for the vaccine failure remain unknown. In the United States alone, CDI results in 20,000 deaths, and the related annual healthcare expenditures exceed $4.8 billion, mainly due to recurrence^8^. Thus, there is a clear need for a next-generation adjuvant that is pharmaceutically acceptable and has a superior immune profile than alum for the vaccine to succeed.

We have established the use of a synthetic liposomal formulation (LS) containing two Toll-like– receptor (TLR) agonists-GLA (TLR4) and 3M-052 (TLR7&8) that work synergistically (GLA 3M-052 LS)^9^. Appropriate selection of the delivery route and adjuvant components allowed the promotion of both mucosal and systemic immune responses that matched the protective immune profile of the target disease. We have also validated the neutralization capacity of both systemic and mucosal antibodies in non-human primates^10^. Based on our experience with the adjuvant platform, we hypothesized that the GLA 3M-052 LS adjuvanted TcdB vaccine will provide a durable, broadly neutralizing, mucosal IgA and systemic IgG immune response to prevent CDI.

In this work, we compared immune responses between inactivated TcdB vaccine adjuvanted either with GLA 3M-052 LS (TcdB-LS) or alum (TcdB-alum) in a murine model of CDI.

In a pilot experiment, we tested the functional ability of plasma samples from the immunized mice to protect target cells upon treatment with TcdB by performing toxin-neutralization assay that evaluates the survival of Chinese Hamster Ovary (CHO) cells following incubation with toxin-plasma mixtures. Mice immunized with TcdB-LS using the intranasal (IN) regimen failed to produce neutralizing plasma IgG response, whereas the intramuscular (IM) regimen showed a poor IgA^+^ memory B response. A combination IM/IN regimen produced more balanced antibody and memory responses (**Supplementary Fig. 1A-C**). Such a combination regimen has previously produced excellent results with a COVID-19 vaccine ^11^. The TcdB-alum group was used as a benchmark based on current clinical trials with *C. difficile* vaccines. In general, alum is not considered suitable for intranasal administration due to potential toxicity concerns. Thus, the TcdB-alum group did not include an IN immunization. All subsequent experiments were carried out using the mixed regimen for TcdB-LS and the IM regimen for TcdB-alum as described. (**Supplementary Fig. 1D**).

For the comparative analysis of TcdB-specific immune responses, three groups of mice (n=8 per group) were immunized three times with a 2-week interval between subsequent immunizations: adjuvant alone (control), TcdB-alum and TcdB-LS. Four weeks after the third immunization, both TcdB-alum and TcdB-LS groups demonstrated equivalent anti-TcdB IgG1 and total IgG titers. Interestingly, the TcdB-LS group exhibited significantly higher levels of IgG2c antibodies **(Fig. 1A)**. Plasma samples were further assessed for avidity. A higher avidity index (AI) indicates better affinity maturation, which may enhance neutralization capabilities. Samples from the TcdB-LS group demonstrated a significantly higher avidity index **(Fig. 1B)**. Anti-TcdB stool and plasma IgA were detected only in the TcdB-LS group **(Fig. 1C and 1D)**. Moreover, the TcdB-LS group also showed significantly higher stool IgG levels **(Fig. 1E)**. A positive correlation was observed between plasma and mucosal IgA levels **(Fig. 1F)**. In conclusion, the liposomal vaccine induced both a balanced systemic humoral and a local stool IgA response.

**Fig 1.**
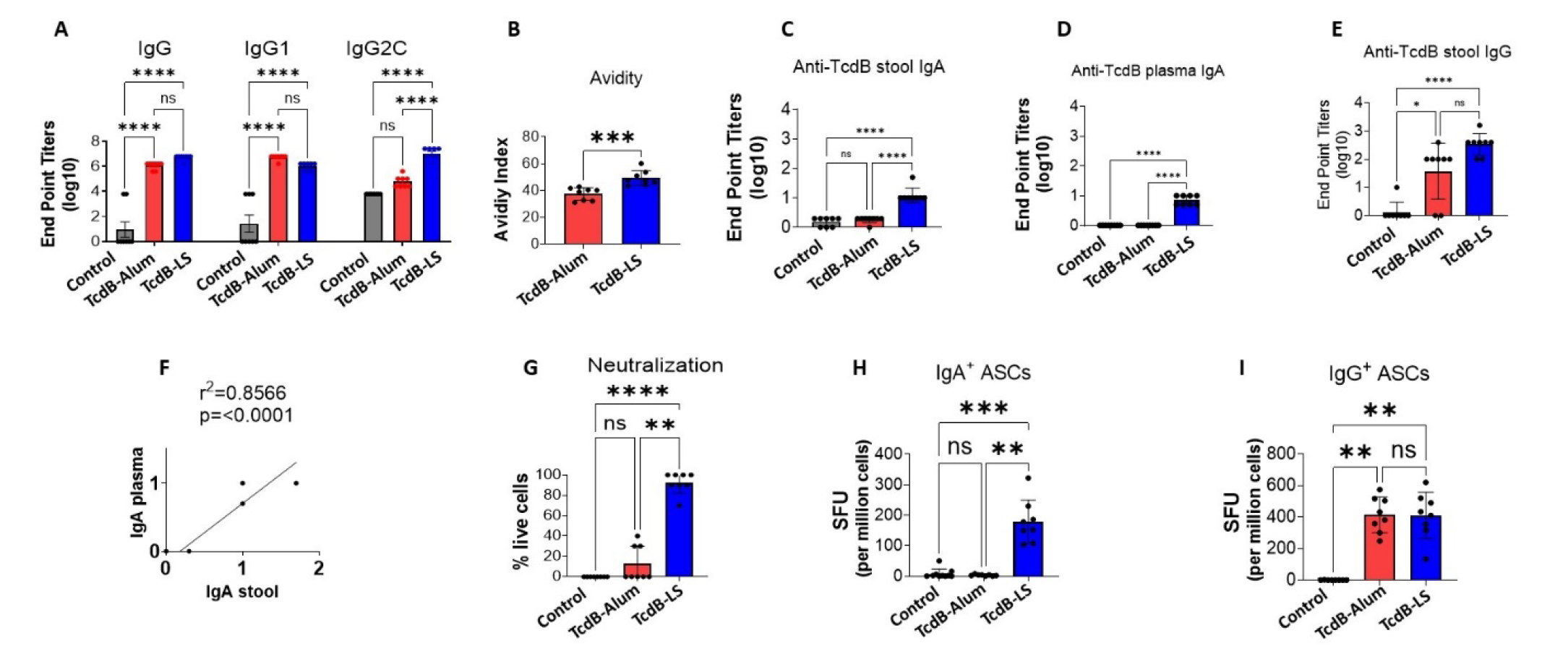
TcdB-LS elicited an enhanced immune response compared to the TcdB-alum: Eight mice per group were immunized three times using TcdB-alum (3 IM) or TcdB-LS (2 IM and an IN boost). Mice receiving only adjuvant served as a control group. Samples were collected four weeks after the final immunization and analyzed. TcdB-B specific IgG, IgG1and IgG2a titers were measured by ELISA using TcdB coated plates. (A) **Plasma anti-TcdB IgG, IgG1 and IgG2c titers**. (B) **Avidity index**. Plasma dilutions that gave an OD450nm = 1.0 were determined and ELISAs for anti-TcdB IgG titers were run as described in two plates in parallel. One plate received standard PBST washes, while the other was washed using 6M urea. Avidity Index (AI %) was expressed: AI = [(absorbance of urea-treated sample/absorbance of non-urea-treated matched sample) × 100]. (C) **Stool anti-TcdB IgA endpoint titers**. (D) **Plasma anti-TcdB IgA endpoint titers**. (E) **Stool IgG endpoint titers**. (F) **Correlation between systemic and mucosal IgA responses in TcdB-LS group**. (G**) Neutralization of TcdB by toxin-specific antibodies**. Various dilutions of plasma from the control and immunized mice were mixed with 4 ng/mL of TcdB and incubated with CHO cells for 24h. Representative data at 1:640 plasma dilution is shown. Complete neutralization of the toxin was characterized by the presence of visually undamaged cells. (H) and (I) **TcdB-specific IgA and IgG antibody-secreting memory cell (ASC) responses**. ELISpot was used to compare antibody-secreting B_MEM_ cells. Single cells were made from the bone marrow 4 weeks post-last immunization and activated with a B cell activator (R848/IL-2) for 72 h. All cells were then cultured in triplicate on TcdB-coated plates for an additional 24 hours. Bone marrow cells from the control mice were used to measure baseline response, which was subtracted from the test readings. Spots were developed, counted, and plotted as spot-forming units (SFU) per million. Two-way ANOVA was used for multiple comparisons. The Mann-Whitney test was used to compare the two groups. Data are represented as mean ± SD. ^*^ p < 0.05, ^**^ p < 0.01, ^***^ p < 0.001, ^****^ p < 0.0001, ns = not significant.

Plasma from the TcdB-LS group demonstrated significantly higher neutralizing antibody activity compared to TcdB-alum **(Fig. 1G)**. Samples from the control group did not exhibit any protection against cell rounding. To assess the TcdB-specific memory B cell (B_MEM_) responses, ELISpot assays were conducted on bone marrow-derived cells. Significantly higher TcdB-specific IgA^+^ antibody-secreting cells (ASCs) were observed in the TcdB-LS group whereas IgG^+^ ASC responses were similar (**Fig. 1H & I)**. Thus, GLA 3M-052 LS supported more robust memory responses.

To compare the protective efficacy between the two adjuvants, immunized mice were challenged with 5 × 10^5^ VPI 10643 spores per mouse, four weeks after the last immunization **(Fig. 2A)**. As expected, control mice succumbed to the challenge and TcdB-LS and TcdB-alum immunized mice were completely protected **(Fig. 2B)**. To assess gut damage between the groups, we conducted a gut-permeability assay 7 days post-challenge. Mice were orally administered FITC-dextran, and gut leakage was determined by measuring its presence in the plasma. The TcdB-LS group showed less gut permeability indicating better protection compared to the TcdB-alum group **(Fig. 2C)**. Bacterial load and the production of toxins were also reduced in both the vaccinated groups **(Supplemental Fig. 2)**.

**Fig 2.**
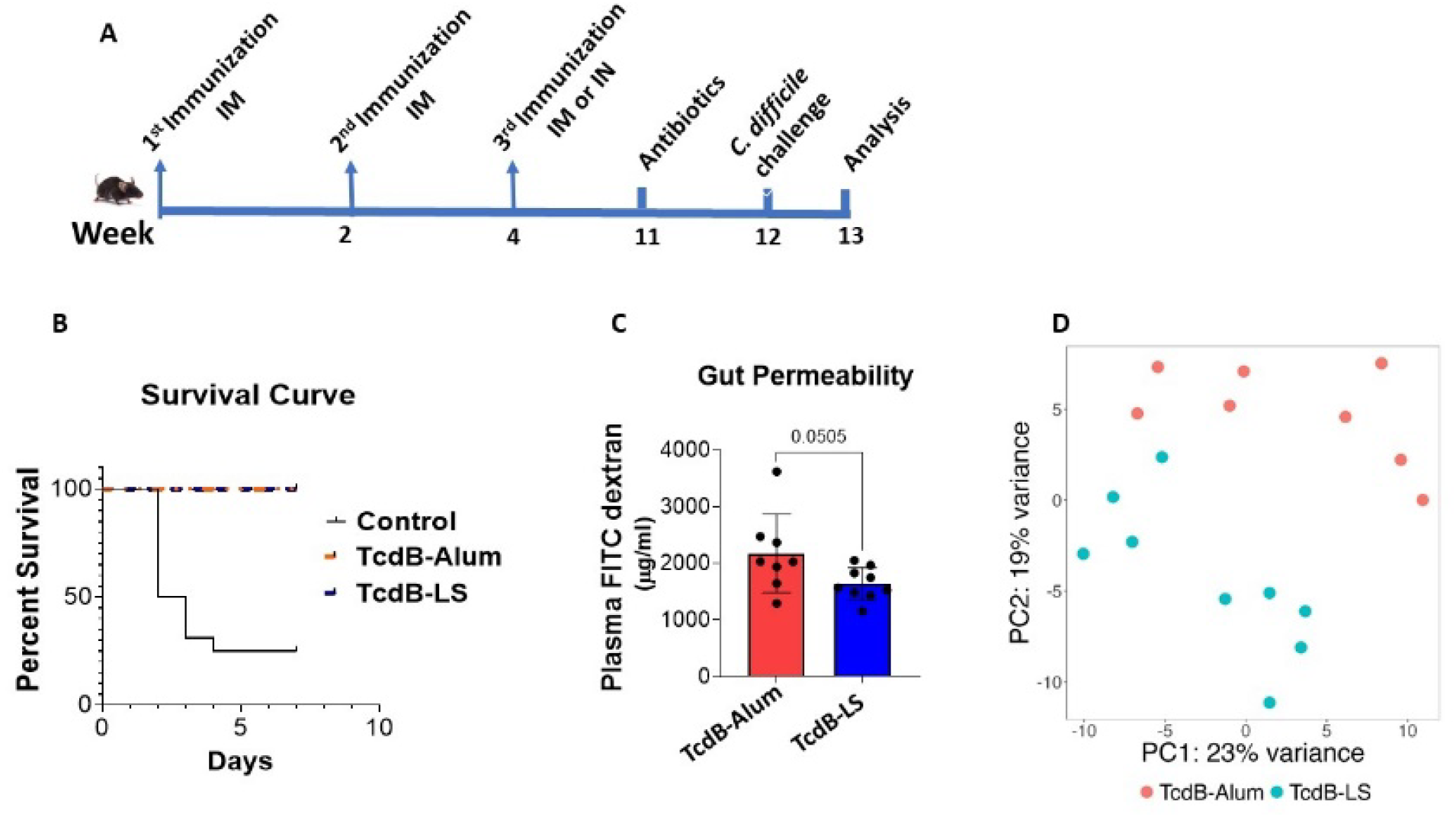
TcdB-LS immunized mice were protected from gut damage during the *C. difficile* infection: 17 Mice per group were challenged with 5 × 10^5^ viable VPI 10643 spores four weeks post-last immunization. Survival and clinical scores were assessed up to 7 days dpi. **(A) Schematic showing immunization and infection timeline. (B) Survival curves. (C) Gut permeability assay**. Mice were fasted for 4 hours at 7 dpi before gavaging with FITC-Dextran. Plasma FITC-dextran concentration leaking in the plasma was measured after an additional 4 hours. **(D) Bulk RNAseq**. Colonic RNA was extracted at 7 dpi. The PCA plot showing transcriptional profiles of TcdB-LS and TcdB-alum groups. The Mann-Whitney test was used to compare the two groups. Data are represented as mean ± SD.

To gain deeper insights into how both the adjuvants modulate the immune response, we conducted bulk RNA sequencing (RNAseq) on colonic tissue from a subgroup of mice (n=8 for TcdB-alum and n=9 for TcdB-LS) 7 days post-infection (dpi). Principal Component Analysis (PCA) illustrated distinct clustering of TcdB-alum and TcdB-LS samples **(Fig. 2D)**, indicating clear differences in their transcriptional profiles. This observation was further validated through differential gene expression analysis, which identified 764 differentially expressed genes (DEGs) (435 upregulated and 329 downregulated in the TcdB-LS group) **(Supplemental Fig. 3)**. Gene Set Enrichment Analysis (GSEA) was employed using Gene Ontology gene sets to identify biological pathways that were significantly enriched within each group^12^. Many immune responses and activation gene sets were preferentially upregulated in the TcdB-LS group compared to the TcdB-alum group **(Table 1)**. Thus, GLA 3M-052 LS promoted a more distinct transcriptional profile as compared to alum.

**Table 1:** Summary of the top enriched immune pathways in the TcdB-LS group compare to the TcdB-alum at 7 dpi as seen from the RNAseq data.

| GO: Biological Processes Gene Set | Normalized Enrichment Score | Adjusted p value | Enriched Genes | Percent Coverage |
| --- | --- | --- | --- | --- |
| Activation of Innate Immune Response | 2.185 | 0.0003 | 36 | 63.158 |
| B Cell Activation | 2.069 | 1.07E-09 | 144 | 44.308 |
| Pattern Recognition Receptor Signaling Pathway | 2.190 | 6.98E-08 | 69 | 40.116 |
| Regulation of CD4 Positive Alpha Beta T Cell Activation | 1.978 | 0.004 | 24 | 38.710 |
| Immunoglobulin Production | 1.784 | 0.002 | 68 | 44.156 |

In the present work, we have utilized our experience with the mucosal adjuvant platform to gain insights into the mechanisms of protection to develop an effective CDI vaccine. The most notable finding was the multifaceted immune profile elicited by the liposomal adjuvant platform given by both IM and IN routes. The molecular and cellular features of the adaptive immune response following CDI are poorly understood. The limited success of Phase 3 clinical trials further highlights the importance of understanding the mechanism of vaccine-mediated protection followed by a rational selection of the adjuvant/route. While GLA and 3M-052 are currently undergoing clinical trials for various conditions, our group is planning to begin a first-in-human Phase 1 trial testing of the combined GLA 3M-052 LS formulation for an amebiasis vaccine candidate.

The pathology associated with CDI is primarily toxin-mediated, and TcdB-specific serum IgG is the only known correlate of protection in humans and animal models^13-16^. GLA 3M-052 LS significantly improved the anti-TcdB IgG response compared to alum, exhibiting higher titers of TcdB-specific IgG2c and better class switching. GLA 3M-052 LS also induced higher avidity IgG antibodies, suggesting a more effective affinity maturation. The neutralization potential of the plasma from the TcdB-LS group was also significantly better when compared to the TcdB-alum group. At this time, we do not know whether the enhanced functional capability could be attributed to higher avidity, efficient class switching, or both.

An ideal vaccine should generate an effective mucosal immune response against a gastrointestinal pathogen and GLA 3M-052 LS has demonstrated its potential to induce long-lasting mucosal responses^11,17^. The role of mucosal IgA in protection against CDI is less clear, but lower fecal IgA concentrations were associated with recurrence^18^. GLA 3M-052 LS elicited an anti-TcdB stool IgA response, whereas alum did not. This enhanced mucosal immunity can provide a critical first line of defense against *C. difficile* to limit tissue damage and shedding of the bacteria.

The high frequency of recurrence in CDI patients suggests a suboptimal memory response following the infection which could affect the quantity and quality of antibodies during the recall response^19,20^. Also, infection failed to restimulate the recall response even in the pre-immunized mice^21^. Therefore, a durable memory response seems to be critical for protection from recurrence. Bone marrow is a key site for the persistence of memory B cells and a long-lasting humoral immunity.^22,23^ ELISpot assay using the bone marrow cells revealed that GLA 3M-052 LS induced a robust B_MEM_ response. In our studies using the GLA 3M-052 LS adjuvanted amebiasis vaccine, we saw a robust and durable systemic as well as mucosal immune response for at least 32 weeks post-final immunization in mice and non-human primates^17^. B_MEM_ cell-encoded IgG1, IgG2b, and IgG2c have the potential to invoke a diverse range of effector functions *in vivo*, and future experiments to determine a role for IgG subclasses in protection are warranted.

Several areas in the *C. difficile* vaccine field need advancement of knowledge, especially after an unexpected termination of clinical trials. These include evaluation of the breadth and neutralizing capacity of the anti-toxin antibodies, the role of mucosal immunity in restricting colonization, and strategies to elicit a robust memory response to avoid frequent recurrence. The liposomal adjuvant platform showed the potential to address several of the knowledge gaps pertaining to gut IgA response, neutralization titer, gut barrier function, and memory.

Both the TcdB-LS and TcdB-alum groups demonstrated 100% protection against *C. difficile* challenge although alum was ineffective in generating mucosal IgA as well as IgA specific B memory response. Alum is known to generate a protective response against a *C. difficile* challenge in mice but not in humans^21,24^. The liposomal group exhibited a better preservation of gut integrity which could be attributed to a more balanced and effective immune response. RNASeq data also supported this observation as seen from the more effective upregulation of key B and T immune pathways in the TcdB-LS group. Overall, GLA 3M-052 LS exhibited a superior target immune profile as compared to alum which may be beneficial in restricting primary as well as recurrent infections.

Several limitations including sex bias, the effect of the immunization route, and antigen dose on the response magnitude and durability still need to be addressed^17^. Cross-protection potential against TcdB subtypes needs to be tested to assess protection from recurrence. Finally, studying transcriptional changes at a time point earlier than 7 dpi may be more informative.

In summary, GLA-3M-052 significantly enhanced systemic and mucosal immune responses, antibody functional capabilities, and memory responses against CDI. The multifaceted immune profile warrants in-depth characterization to address knowledge gaps and work toward an effective CDI vaccine.

## Methods

### Antigen and adjuvant

Inactivated toxin B (toxoided TcdB) was acquired from TechLab, Inc. (Blacksburg, VA). Alum (Alhydrogel 2%, Croda) was diluted in water by AAHI and supplied for the immunization studies. GLA 3M-052 LS has been described previously and was also supplied by AAHI ^9^.

### Animals and Immunizations

Animal procedures were conducted in accordance with the guidelines approved by the Institutional Animal Care and Use Committee at the University of Virginia (IACUC). Eight-week-old C57BL/6J mice were procured from room EMO3 of the Jackson Laboratory and distributed randomly across the groups. All the treatments (immunizations, infection) were also randomly allocated to ensure equal representation across the groups. Toxoided TcdB (20 μg) was mixed with either Alum (100 µg) or Liposome adjuvant (10 µg GLA, 4 µg 3M-O52) formulations just before immunizations. Volume was adjusted to 100 μL for intramuscular and

50 μL for intranasal immunization using saline (n=8 per group). Intramuscular injections targeted the thigh region. Intranasal immunizations were performed with 25 μL of inactivated

TcdB-adjuvant mix administered per nostril after the animals were anesthetized using an intraperitoneal Ketamine/ Xylazine (60-80/5-10 mg/kg) injection. Animals were observed until fully awake after intranasal immunizations and all efforts were made to minimize animal suffering. A two-week interval was maintained between successive immunizations across all regimens, totaling three immunizations. Immunogenicity was evaluated four weeks after the final immunization. Animals were euthanized using an overdose of anesthesia (twice the anesthetic dose) via intraperitoneal route followed by cervical dislocation.

### Fecal toxin detection

Fecal toxin A/B in the cecal contents was detected using the C. DIFFICILE TOX A/B II ELISA kit (TechLab Inc., catalog #T5015). *C. difficile* glutamate dehydrogenase was measured using the C. DIFF CHEK -60 kit (TechLab Inc., catalog #TL5025). A standard curve was used to represent bacterial load as arbitrary units.

### ELISA

Plasma, cecal contents, and stool samples were stored at -80°C until they were analyzed for TcdB-specific IgG and IgA antibody titers using an enzyme-linked immunosorbent assay (ELISA). 96-well ELISA half-area plates (Corning Costar, Cat# 3690) were coated with 2 µg/well of toxin B (Techlab) in 50 mM bicarbonate buffer (pH 9.3, Sigma-C3041) overnight at 4°C. After coating, the plates were washed three times with 1x PBS-Tween 20 (PBST; Thermo Scientific, Cat# 28352) using a BioTek plate washer and then blocked with commercial ELISA blocking buffer (Thermo Scientific, Cat# N502) for 1 hour at room temperature. Subsequently, the plates were air-dried and stored until further use.

Prior to mucosal antibody analysis, all stool and cecal content samples were processed as follows: to each milligram of stool or cecal content, 10 µL of PBS containing protease inhibitors (Thermo Scientific, Cat# A32953) was added. The samples were homogenized using a bead beater (Qiagen, Germantown, MD) at 250 rev/min for 10 minutes and then centrifuged at 14,000 x g at 4°C for 10 minutes. The resulting supernatants were carefully transferred into 1.5-mL microcentrifuge tubes (Axygen), and phenylmethylsulfonyl fluoride was added to a final concentration of 1 mM before storage at -80°C.

Thawed plasma samples were used directly without additional processing. Serial dilutions of plasma were prepared with ELISA blocking buffer and added to the toxin B-coated wells of the ELISA plates, followed by incubation for 1 hour at 37°C. After washing with PBST, horseradish peroxidase (HRP)-conjugated antibodies (Goat Anti-Mouse IgG1 at 1:5000, Goat Anti-Mouse IgG2c at 1:1000, Goat Anti-Mouse IgG at 1:5000, and Goat Anti-Mouse IgA at 1:1000, sourced from SouthernBiotech and catalog #1040-05) were added and incubated for 1 hour at 37°C. The color was developed using 1-Step™ Ultra TMB-ELISA Substrate Solution (Thermo Scientific 34028), and the reaction was stopped with 1:100 diluted sulfuric acid (Sigma-Aldrich). Optical densities (ODs) were measured at 450 nm using an ELISA reader (Synergy, BioTek). Endpoint titers were determined with OD cutoffs of 0.5 for plasma IgG and 0.1 for mucosal IgA.

### IgG avidity assay

The avidity index (AI) was measured as previously described ^25^. The avidity of antibodies specific to toxin B was assessed using a urea disruption ELISA. To summarize, ELISA plates were coated with 2 µg/mL of toxin B. Plasma samples were diluted to achieve a value of 1.0 at OD_450nm_ ensuring that the antibody concentrations were within a linear range. Samples were plated in triplicate on two sets of ELISA plates: one set washed with standard PBST and the other with a dissociation buffer (PBS containing 6 M urea). Both sets were then incubated with an HRP-conjugated secondary antibody, developed, and read. The avidity percentage was determined using the following formula: Avidity Index (AI) = [(average absorbance of urea-treated sample/average absorbance of PBST-treated matched sample) × 100].

### In-vitro neutralization assay

The neutralization of TcdB by plasma antibodies was assessed using Chinese hamster ovary (CHO) cell lines. CHO cells were seeded at a density of 3 × 10^4^ cells per well in a 96-well microtiter plate and incubated for 24 hours at 37°C with 5% CO2 in Ham’s F-12K medium (Thermo Fisher #21127022) supplemented with 10% heat-inactivated fetal bovine serum (HI FBS) and 1% penicillin-streptomycin, to reach 70-80% confluence. Toxin B was mixed with the plasma from immunized mice at various concentrations in PBS and incubated for 60 minutes at 37°C. The CHO cells were washed with PBS, resuspended in 75 µL of media, overlaid with 25 µL of the plasma-toxin mix, and incubated at 37°C for 24 hours in 5% CO2. The cytotoxic effect of non-neutralized toxins was evaluated microscopically, indicated by the presence of cells exhibiting partial to complete rounding. Complete neutralization of the toxin was characterized by the presence of visually undamaged cells. Toxin neutralization by sera from immunized mice was tested using twofold dilutions (ranging from 1/10 to 1/80) incubated with 2 ng/mL of toxin B.

### ELISpot assays

B cell ELISpot assays were performed to quantify the memory B cell (B_MEM_) pool, which is essential for generating a secondary immune response upon antigen re-exposure. Ninety-six-well multi-screen filter plates (Millipore, Cat# MSIPS4W10) were pre-treated with 35% ethanol and subsequently rinsed with sterile PBS. The wells were then coated with 2 µg/mL of TcdB to evaluate vaccine-specific responses. As controls, additional plates were also coated with 15 µg/mL of anti-mouse IgG or anti-mouse IgA capture antibodies to measure total IgG and IgA-secreting cells. Bone marrow cells were harvested from the femurs of immunized mice and cultured in RPMI with 10% FBS, and activated with the B-Poly-S Polyclonal B Cell Activator (R848/IL-2) for 48 hours before spot development. Approximately 500,000 activated cells were seeded per well in 200 µL of medium and incubated at 37°C in a CO2 incubator for 48 hours. An AEC substrate solution was added for up to 15 minutes to visualize the spots, and the reaction was terminated by rinsing under running distilled water. The plates were dried in the dark for a minimum of 2 days, and the spots were counted using an ELISpot reader. The reported values were averaged from two dilutions. Reference plasma from toxoid B-immunized mice served as a positive control, while plasma from unimmunized mice served as a negative control.

### Bacterial strains and culture

Mice were infected with 10^5^ CFU/mL of *Clostridioides difficile* VPI 10643 strain spores. *C. difficile* strains were cultured on Brain Heart Infusion (BHI) agar from glycerol stocks and incubated overnight at 37°C under anaerobic conditions. Columbia, clospore, and BHI broths were pre-reduced for a minimum of 24 hours. VPI spore stocks were prepared according to a previously described protocol ^26^. Briefly, a single colony was inoculated into 15 mL of Columbia broth and incubated overnight at 37°C. Subsequently, 5 mL of this culture was anaerobically transferred to 45 mL of Clospore broth and incubated for 7 days at 37°C. The resulting culture was washed at least five times with cold, sterile water and resuspended in 1 mL of sterile water. Spores were stored in 1.5 mL twist-cap tubes at 4°C (Corning #4309309). Each mouse received 100 μL of the inoculum via oral gavage. The actual inoculum concentration was verified by plating on BHI agar supplemented with 0.032 mg/mL cefoxitin, 1 mg/mL D-cycloserine, and 1% sodium taurocholate (Sigma), followed by anaerobic incubation at 37°C overnight.

### Infection Model

Mice were infected with *C. difficile* as previously described ^27^. Briefly, mice (n=17 per group) received an antibiotic cocktail in their drinking water consisting of 215 mg/L metronidazole (Hospira), 35 mg/L colistin (Sigma), 45 mg/L vancomycin (Mylan), and 35 mg/L gentamicin (Sigma) for 3 days, followed by regular drinking water. One day prior to infection, clindamycin (Hospira) was administered intraperitoneally at a dose of 0.016 mg/g. Post-infection, mice were monitored twice daily to assess clinical scoring parameters and weight loss. The scoring criteria included weight loss, coat condition, eye condition, activity level, diarrhea, and posture, as described ^27^. Mice were humanely euthanized using an overdose of anesthesia followed by cervical dislocation as described if they reached a clinical score of 14 or experienced more than 25% weight loss. The investigators were not blinded during the studies.

### FITC-dextran gut permeability assay

After fasting for four hours, mice were gavaged with fluorescein isothiocyanate (FITC)-dextran solution (Sigma-Aldrich, # 46944-500MG-F) at a dose of 44 mg per 100 g of body weight. Four hours post-gavage, the mice were euthanized, and plasma samples were collected. The FITC-dextran present in the plasma was measured using a spectrophotometer at wavelengths of 485/530 nm.

### RNA Isolation, Sequencing, and Analysis

RNAlater-stabilized cecal tissue samples were excised and subsequently immersed in 1mL of TRIzol per tissue sample. Homogenization of the tissues was carried out using buffer RLT with 1% β-mercaptoethanol (Qiagen). The aqueous phase was isolated and purified utilizing the RNeasy Plus mini kit (Qiagen) following the manufacturer’s instructions. RNA sequencing was performed by Novogene, employing polyA enrichment and sequencing on an Illumina NovaSeq X Plus platform with paired-end 150 bp reads. Quality assessment of unprocessed FASTQ files was conducted using FastQC (v0.12.1)^28^ and MultiQC (v1.14)^29^. The reads were pseudo-aligned to the murine genome (v109 from EnsemblDB) using Kallisto (v0.44.0)^30^. Count tables were imported into R with TxImport (v1.30.0)^31^, and the DESeq2 package (v1.42.0)^32^ was utilized to filter low-count genes, normalize the data, estimate dispersions, and fit counts using a negative binomial model. Differential gene expression analysis was conducted, ranking genes according to Wald’s statistic from most upregulated to most downregulated post-FMT. This ranked list was used for Gene Set Enrichment Analysis (GSEA) with the fgsea package (v1.20.0)^33^ using Gene Ontology databases ^34^.

### Statistical analyses

Statistical analyses were performed using Graph Pad Prism software and Microsoft Excel (www.biostathandbook.com/welchanova.xls). Statistically significant differences were determined by one-way ANOVA with appropriate correction for multiple comparisons. No animals were excluded during the analysis.

## Data availability

Unprocessed FASTQ files from the RNASeq data generated in this study have been deposited in the NCBI Sequence Read Archive (SRA) under Accession number PRJNA1139815. All the codes used for RNAseq analysis will be deposited in GitHub at https://github.com/petrilab-uva/2024-CDI-vaccine.

## Acknowledgments

The authors gratefully acknowledge Robert Kinsey and Gabi Ramer-Denisoff from AAHI for preparing and characterizing the adjuvants.

## Author Contributions

FN: Data curation, formal analysis, investigation, methodology, validation, visualization, writing original draft, review, and editing. NH: methodology, validation, review, and editing. BT: methodology. GBM: RNA sequencing data analysis. MY: methodology. JH: Material, review. CBF: Material, review. FX: methodology. MMA: Conceptualization, formal analysis, funding acquisition, methodology, project administration, supervision, review, and editing. WAP: Conceptualization, formal analysis, funding acquisition, methodology, project administration, supervision, review, and editing.

## References

1 Ofosu, A. Clostridium difficile infection: a review of current and emerging therapies. Ann Gastroenterol 29, 147–154 (2016). 10.20524/aog.2016.0006

2 Marra, A. R. et al. Incidence and Outcomes Associated With Clostridium difficile Infections: A Systematic Review and Meta-analysis. Jama Netw Open 3 (2020). https://doi.org:ARTNp e1917597 10.1001/jamanetworkopen.2019.17597

3 Rupnik, M., Wilcox, M. H. & Gerding, D. N. Clostridium difficile infection: new developments in epidemiology and pathogenesis. Nat Rev Microbiol 7, 526–536 (2009). 10.1038/nrmicro2164

4 Shen, E. H. et al. Subtyping analysis reveals new variants and accelerated evolution of Clostridioides difficile toxin B. Commun Biol 3 (2020). 10.1038/s42003-020-1078-y

5 King, A. M., Mackin, K. E. & Lyras, D. Emergence of toxin A-negative, toxin B-positive Clostridium difficile strains: epidemiological and clinical considerations. Future Microbiol 10, 1–4 (2015). 10.2217/fmb.14.115

6 de Bruyn, G. et al. Safety, immunogenicity, and efficacy of a Clostridioides difficile toxoid vaccine candidate: a phase 3 multicentre, observer-blind, randomised, controlled trial. Lancet Infect Dis 21, 252–262 (2021). 10.1016/S1473-3099(20)30331-5

7 Christensen, S. et al. A phase 3 study evaluating the lot consistency, immunogenicity, safety, and tolerability of a Clostridioides difficile vaccine in healthy adults 65 to 85 years of age. Vaccine 41, 7548–7559 (2023). 10.1016/j.vaccine.2023.11.003

8 Naz, F. & Petri, W. A. Host Immunity and Immunization Strategies for Clostridioides difficile Infection. Clin Microbiol Rev 36, e0015722 (2023). 10.1128/cmr.00157-22

9 Abhyankar, M. M. et al. Adjuvant composition and delivery route shape immune response quality and protective efficacy of a recombinant vaccine for Entamoeba histolytica. NPJ Vaccines 3, 22 (2018). 10.1038/s41541-018-0060-x

10 Abhyankar, M. M. et al. Immunogenicity and safety of an Entamoeba histolytica adjuvanted protein vaccine candidate (LecA+GLA-3M-052 liposomes) in rhesus macaques. Hum Vaccin Immunother 20, 2374147 (2024). 10.1080/21645515.2024.2374147

11 Abhyankar, M. M. et al. Development of COVID-19 vaccine using a dual Toll-like receptor ligand liposome adjuvant. NPJ Vaccines 6, 137 (2021). 10.1038/s41541-021-00399-0

12 Subramanian, A. et al. Gene set enrichment analysis: a knowledge-based approach for interpreting genome-wide expression profiles. Proc Natl Acad Sci U S A 102, 15545–15550 (2005). 10.1073/pnas.0506580102

13 Ananthakrishnan, A. N. Clostridium difficile infection: epidemiology, risk factors and management. Nat Rev Gastroenterol Hepatol 8, 17–26 (2011). 10.1038/nrgastro.2010.190

14 Donald, R. G. K. et al. A novel approach to generate a recombinant toxoid vaccine against Clostridium difficile. Microbiology (Reading) 159, 1254–1266 (2013). 10.1099/mic.0.066712-0

15 Steele, J., Mukherjee, J., Parry, N. & Tzipori, S. Antibody against TcdB, but not TcdA, prevents development of gastrointestinal and systemic Clostridium difficile disease. J Infect Dis 207, 323–330 (2013). 10.1093/infdis/jis669

16 Wilcox, M. H. et al. Bezlotoxumab for Prevention of Recurrent Clostridium difficile Infection. N Engl J Med 376, 305–317 (2017). 10.1056/NEJMoa1602615

17 Abhyankar, M. M. et al. Optimizing a Multi-Component Intranasal Entamoeba Histolytica Vaccine Formulation Using a Design of Experiments Strategy. Front Immunol 12, 683157 (2021). 10.3389/fimmu.2021.683157

18 Johal, S. S. et al. Colonic IgA producing cells and macrophages are reduced in recurrent and non-recurrent Clostridium difficile associated diarrhoea. J Clin Pathol 57, 973–979 (2004). 10.1136/jcp.2003.015875

19 Shah, H. B. et al. Human C. difficile toxin-specific memory B cell repertoires encode poorly neutralizing antibodies. Jci Insight 5 (2020). https://doi.org:ARTN e138137 10.1172/jci.insight.138137

20 Norman, K. M. et al. Clostridioides difficile toxin B subverts germinal center and antibody recall responses by stimulating a drug-treatable CXCR4-dependent mechanism. Cell Rep 43, 114245 (2024). 10.1016/j.celrep.2024.114245

21 Amani, S. A., Shadid, T., Ballard, J. D. & Lang, M. L. Clostridioides difficile Infection Induces an Inferior IgG Response to That Induced by Immunization and Is Associated with a Lack of T Follicular Helper Cell and Memory B Cell Expansion. Infect Immun 88 (2020). https://doi.org:ARTN e00829–19 10.1128/IAI.00829-19

22 Palm, A. E. & Henry, C. Remembrance of Things Past: Long-Term B Cell Memory After Infection and Vaccination. Front Immunol 10, 1787 (2019). 10.3389/fimmu.2019.01787

23 Chang, H. D. & Radbruch, A. Maintenance of quiescent immune memory in the bone marrow. Eur J Immunol 51, 1592–1601 (2021). 10.1002/eji.202049012

24 Devera, T. S. et al. Memory B Cells Encode Neutralizing Antibody Specific for Toxin B from the Clostridium difficile Strains VPI 10463 and NAP1/BI/027 but with Superior Neutralization of VPI 10463 Toxin B. Infect Immun 84, 194–204 (2016). 10.1128/Iai.00011-15

25 Beghetto, E. et al. Use of an immunoglobulin G avidity assay based on recombinant antigens for diagnosis of primary Toxoplasma gondii infection during pregnancy. J Clin Microbiol 41, 5414–5418 (2003). 10.1128/JCM.41.12.5414-5418.2003

26 Donlan, A. N., Simpson, M. E. & Petri, W. A., Jr. Type 2 cytokines IL-4 and IL-5 reduce severe outcomes from Clostridiodes difficile infection. Anaerobe 66, 102275 (2020). 10.1016/j.anaerobe.2020.102275

27 Frisbee, A. L. et al. IL-33 drives group 2 innate lymphoid cell-mediated protection during Clostridium difficile infection. Nat Commun 10, 2712 (2019). 10.1038/s41467-019-10733-9

28 Andrews, S. (Cambridge, United Kingdom, 2010).

29 Ewels, P., Magnusson, M., Lundin, S. & Kaller, M. MultiQC: summarize analysis results for multiple tools and samples in a single report. Bioinformatics 32, 3047–3048 (2016). 10.1093/bioinformatics/btw354

30 Bray, N. L., Pimentel, H., Melsted, P. & Pachter, L. Near-optimal probabilistic RNA-seq quantification. Nat Biotechnol 34, 525–527 (2016). 10.1038/nbt.3519

31 Soneson, C., Love, M. I. & Robinson, M. D. Differential analyses for RNA-seq: transcript-level estimates improve gene-level inferences. F1000Res 4, 1521 (2015). 10.12688/f1000research.7563.2

32 Love, M. I., Huber, W. & Anders, S. Moderated estimation of fold change and dispersion for RNA-seq data with DESeq2. Genome Biol 15, 550 (2014). 10.1186/s13059-014-0550-8

33 Korotkevich, G. et al. Fast gene set enrichment analysis. bioRxiv, 060012 (2021). 10.1101/060012

34 Ashburner, M. et al. Gene ontology: tool for the unification of biology. The Gene Ontology Consortium. Nat Genet 25, 25–29 (2000). 10.1038/75556

