## Supplemental figures for "Enhanced immunogenicity of a *Clostridioides difficile* TcdB vaccine adjuvanted with a synthetic dual-TLR ligand adjuvant"

**week**

**2**

**4**

**8**

**Analysis**

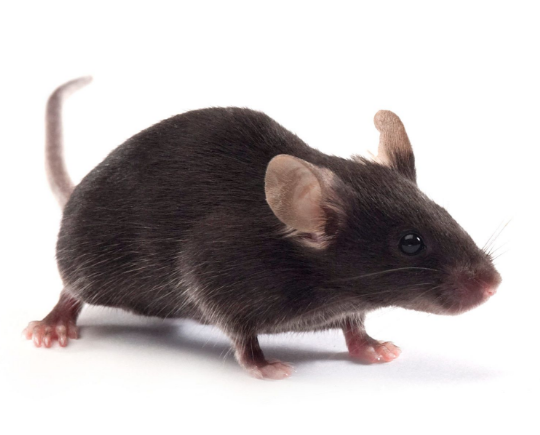

**1^st^ Immunization**

**IM**

**3^rd^ Immunization**

**IM or IN**

**2^nd^ Immunization**

**IM**

**Supplementary Fig 1. (A) Neutralization assay.** Various dilutions of plasma from the control and immunized mice were mixed with 4 ng/mL of TcdB and incubated with CHO cells for 24h. Complete neutralization of the toxin was characterized by the presence of visually undamaged cells. Representative data at 1:640 plasma dilution is shown. **(B)** IgA+ memory B assay. TcdB-specific IgA antibody-secreting cell (ASC) responses in bone marrow were assessed by ELISpot. Single cells were made from the bone marrow 4 weeks post-last immunization and activated with a B cell activator (R848/IL-2) for 72 h. All cells were then cultured in triplicate on TcdB-coated plates for an additional 24 hours. Bone marrow cells from the control mice were used to measure baseline response, which was subtracted from the test readings. Spots were developed, counted, and plotted as spot-forming units (SFU) per million. **(C)** Anti-TcdB specific total IgG titers. Plasma samples were collected four weeks after the final immunization and analyzed by ELISA. **(D)** Schematic showing immunization timeline. * p < 0.05, ** p < 0.01, ***p< 0.005, **** p< 0.001, ns = not significant.

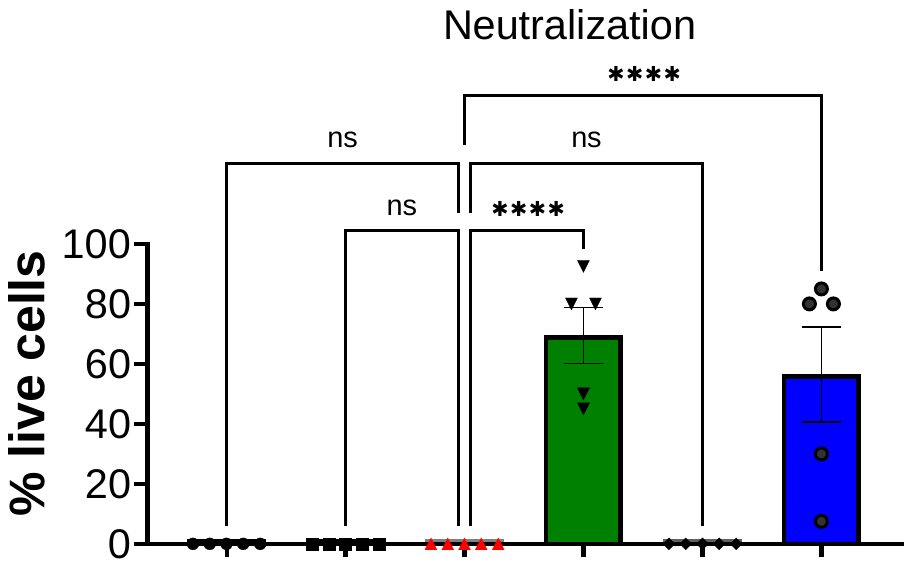

**Saline only**

**TcdB-Alum (IM)**

**TcdB-LS (IM/IN)**

**Antigen only (IM)**

**TcdB-LS (IM)**

**TcdB-LS (IN)**

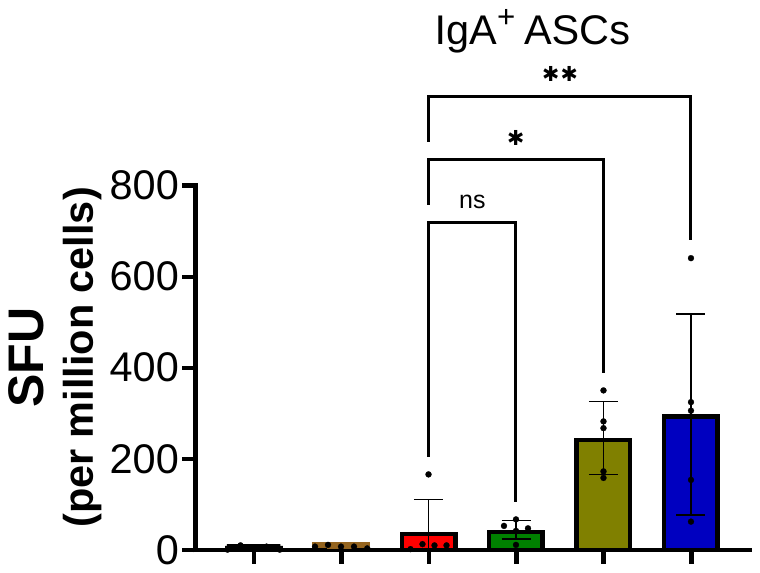

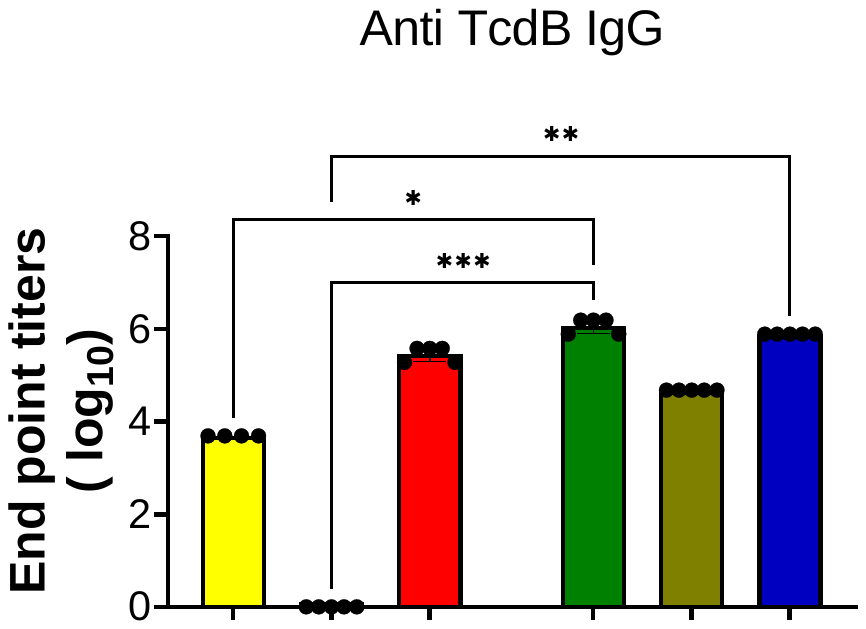

**A**

**B**

**C**

**Saline only**

**TcdB-Alum (IM)**

**TcdB-LS (IM/IN)**

**Antigen only (IM)**

**TcdB-LS (IM)**

**TcdB-LS (IN)**

**Saline only**

**TcdB-Alum (IM)**

**TcdB-LS (IM/IN)**

**Antigen only (IM)**

**TcdB-LS (IM)**

**TcdB-LS (IN)**

**Supplemental Figure 1**

**D**

**Supplemental Figure 2**

**B**

**A**

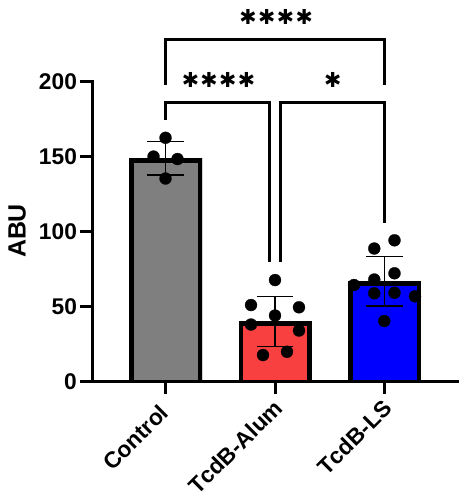

**Toxin A/B ELISA**

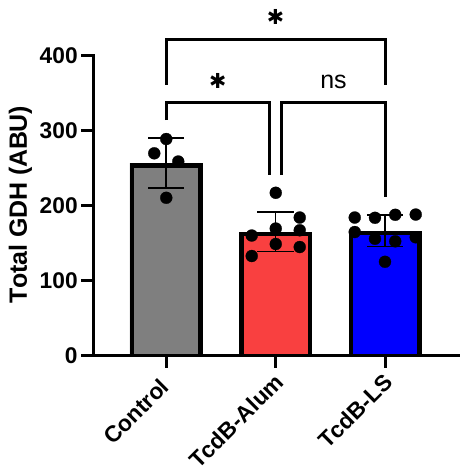

**Relative bacterial abundance**

**Supplementary Fig 2.** Vaccination reduced bacterial burden and toxin production in mice **(A)** The C. DIFF CHEK - 60 kit (TechLab Inc., catalog #TL5025) was used according to the manufacturer’s instructions to quantify bacterial count. **(B)**The TOX A/B II ELISA kit (TechLab Inc., catalog #T5015) was used according to the manufacturer’s instructions to quantify Toxin A/B. Data was normalized to the weight of cecal content. One-way ANOVA was used for the analysis. * p < 0.05, ****p< 0.0001, ns = not significant. ABU= arbitrary units.

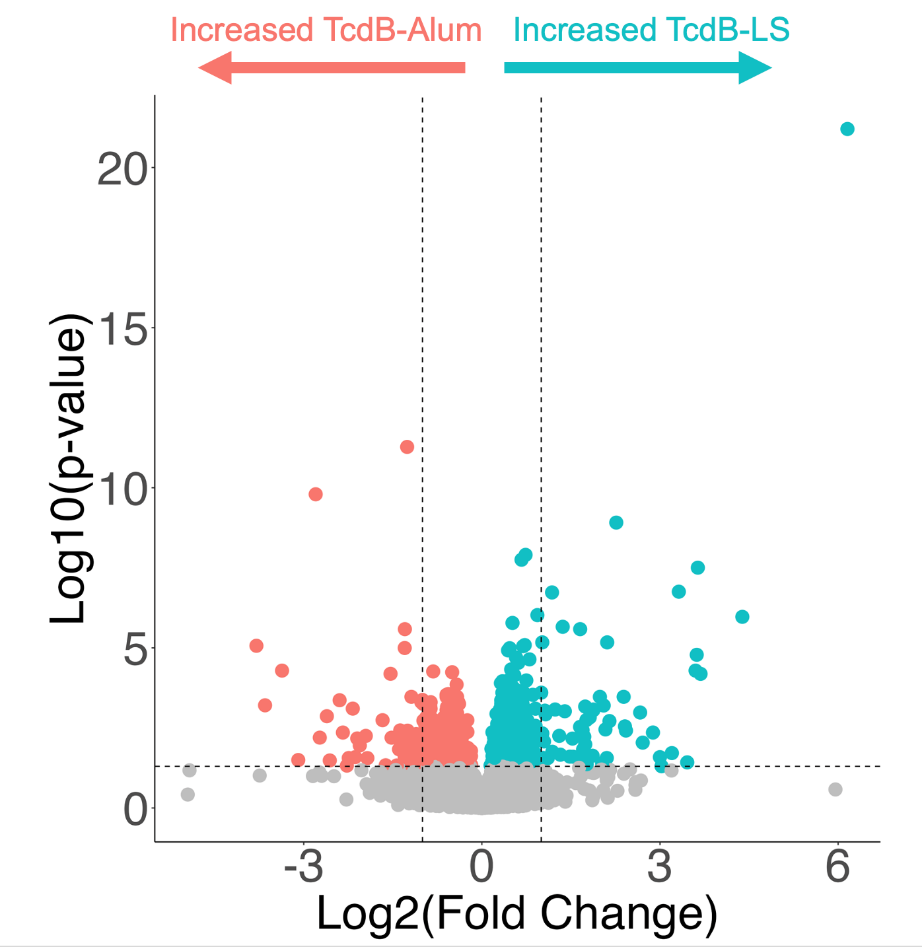

**Supplementary Fig 3.** Alum and liposome adjuvant displayed distinct transcriptional profiles post challenge: Volcano plot analysis representing differentially expressed genes (DEGs) between the groups. The horizontal dotted line represents an adjusted p-value cutoff of 0.05, while vertical dotted lines represent 2-fold change cutoffs. DEGs were calculated from a DESeq2 multivariate model using Wald’s test with FDR correction for multiple comparisons.

**Supplemental Figure 3**
